# Microbial communities on station and train surfaces in Chennai Metro: Insights into urban transit microbiome

**DOI:** 10.1101/2025.03.12.642787

**Authors:** Vijaya Yuvaram Singh, Veerendra P. Gadekar, Srijith Sasikumar, R. M. Rajeeva Lokshanan, V. Senthamizhan, Bharath Prithiviraj, Himanshu Sinha, Karthik Raman

**Author notes:** Corresponding authors: Karthik Raman, Himanshu Sinha. These authors contributed equally to the work (Vijaya Yuvaram Singh, Veerendra P. Gadekar).

## Abstract

**Background:** Urban public transport systems, particularly metro networks, serve as key hubs for microbial transmission, yet the urban microbiome in densely populated regions like India remains poorly characterized. These environments harbour diverse microbial communities, including beneficial and pathogenic species, which can influence public health. The COVID-19 pandemic has further underscored the need to monitor microbial ecosystems, particularly with respect to antimicrobial resistance (AMR) genes, which may have escalated due to the increased use of antibiotics during health crises. In a first-of-its-kind study in India, we comprehensively characterized microbial communities and the prevalence of AMR genes in the Chennai Metro system. We collected 96 surface swab samples from 12 stations across two metro lines, targeting four surfaces: bannisters, kiosks, rods, and ticket counters. Forty-seven samples passed quality control and were analyzed using whole-genome metagenomic sequencing. We employed taxonomic classification and conducted comparative and diversity analyses, as well as differential taxa profile analyses, across the sample collection objects. We studied the prevalence and abundance of AMR genes using AMR annotations from the CARD database. We performed pangenome analysis by constructing Metagenome Assembled Genomes (MAGs) from the collected samples and comparing them with the NCBI reference genome.

**Results:** Comparative analysis with global urban microbiome datasets revealed distinct microbial profiles, including nine species that are differentially prevalent in Chennai samples. Surface type significantly influenced microbial diversity, with kiosks exhibiting the highest diversity. We successfully reconstructed several high-quality MAGs, providing insights into the genomic potential and adaptability of dominant taxa in this environment. While the overall prevalence of AMR genes was minimal, genes associated with Sulfonamide and Rifamycin resistance were detected.

**Conclusion:** These findings highlight unique microbial signatures and emphasize the need for ongoing surveillance and targeted interventions to mitigate microbial transmission risks in densely populated urban areas.

## INTRODUCTION

Public transport systems represent a dynamic environment for microbial transmission, facilitating daily interactions among diverse populations and their microbiomes. The metro system, in particular, acts as a hub for a wide range of microorganisms, including beneficial commensals and symbionts, as well as potentially pathogenic bacteria, making it a potential vector for the spread of infectious diseases [1, 2]. As critical infrastructure in urban settings, public transportation systems significantly influence the microbial ecology of cities. Global initiatives, such as the MetaSUB consortium [3, 4], have advanced our understanding of urban microbiomes by characterizing the unique microbial profiles of cities around the world, providing crucial data for urban planning and public health interventions. Despite the increasing number of studies on urban microbiomes, densely populated Indian cities remain largely unexplored. As one of the fastest-growing urban centers in the world, Chennai, with a population exceeding 12 million [5], exemplifies the dramatic transformation of Indian cities in the 21st century. This study represents the first comprehensive analysis of urban microbiomes in public transportation systems in India, offering critical insights into the unseen biological ecosystems that thrive in these densely populated spaces.

Understanding the microbial communities in public transit systems is crucial for urban public health and global AMR surveillance. AMR, often described as a hidden pandemic [6], has intensified due to rising antibiotic use during health crises such as the COVID-19 pandemic. In a metropolis like Chennai, where environmental factors, human activity, and rapid urbanization intersect, the dynamics of microbial communities, including their potential to harbour AMR genes, are still poorly understood. Unlike subway systems in extensively studied cities, Chennai’s unique socio-environmental landscape presents a distinct microbial signature, making it a key region for global comparative microbiome studies.

Recent advancements in next-generation sequencing (NGS) technologies have revolutionized our understanding of microbial communities, particularly those that are difficult to cultivate in the laboratory. The increasing availability of NGS data in public repositories has facilitated unprecedented opportunities to track the dissemination of microorganisms within urban environments [7–9]. Investigating bacterial communities in regions severely affected by the COVID-19 pandemic is crucial for understanding the dynamics of AMR gene dissemination, which may have been amplified by the increased use of antibiotics during the pandemic [10, 11].

Urban microbiome research has experienced significant growth in recent years, driven by initiatives such as the MetaSUB consortium, which aims to map microbial ecosystems in cities worldwide. Yet, despite its vast and densely populated urban centers, India remains largely unexplored in this context. This study addresses that gap by characterizing the Chennai metro’s microbiome, uncovering distinct microbial signatures, including nine core species that have not been previously identified in other global cities. These findings underscore the need to examine how urban environments in developing nations shape microbial diversity and antimicrobial resistance patterns.

By characterizing the microbial landscape of Chennai’s Metro system, we provide the first window into how such environments in India may serve as reservoirs for both beneficial and pathogenic microorganisms. Moreover, this work is a call to action to implement continuous surveillance systems in urban transit environments, where rapid microbial transmission could pose heightened risks during future public health crises.

## RESULTS

### Taxonomic classification and comparative analysis of distinct microbial signatures

We first investigated how microbial species are distributed across the environments of Chennai metro stations (Figure 1). Specifically, we aimed to determine whether the urban environment exhibited a homogeneous microbial ecosystem or comprised distinct yet interconnected communities, particularly with respect to biodiversity. To achieve this, we examined the prevalence of species, defined as the fraction of samples in which a given taxon was found at different thresholds of relative abundance. We identified a right-skewed distribution characterized by a prominent primary peak on the left, representing the majority of species with low prevalence. A smaller secondary peak on the right highlighted a subset of species with distinct characteristics or higher prevalence within the sampled ecosystem

**Figure 1:**
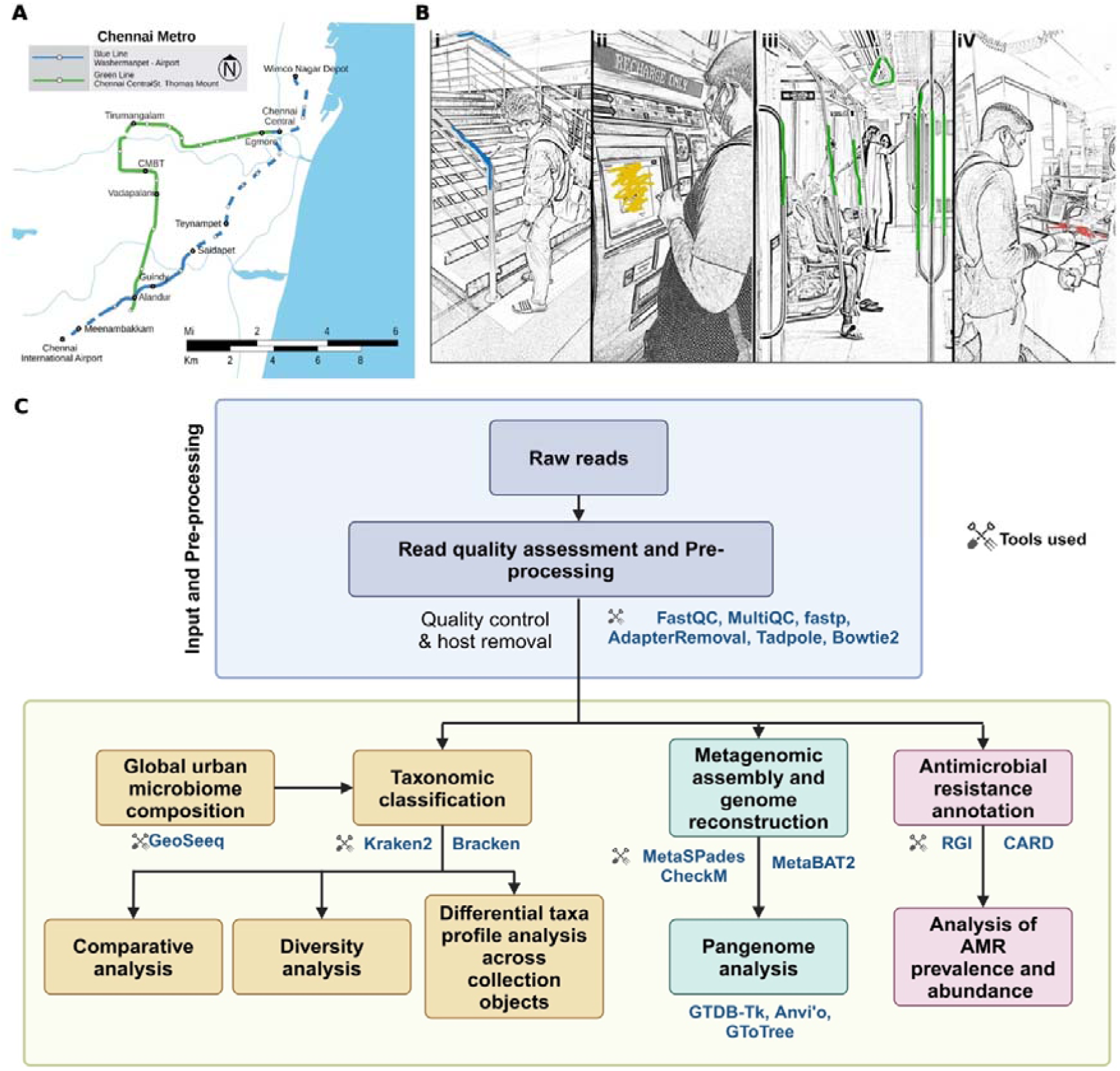
(A) Chennai Metro Line Map. Highlighting sample collection stations [47] (B) Samples were collected from various surfaces, including – (i) handrails in metro stations (bannisters), (ii) ticket vending machines (kiosks), (iii) handrails in trains (rods), and (iv) ticket counters. The swabbing areas on these surfaces are highlighted in blue, yellow, green, and red, respectively. To maintain consistency across all samples, only a portion of each surface was swabbed rather than the entire surface. (C) Workflow for data processing and analysis.

(Figure 2A). This allowed us to categorize the taxa into three distinct groups: ‘core’ taxa, present in the majority of samples; ‘sub-core’ taxa, present in 80-97% of samples; and ‘peripheral’ species, observed in fewer than 15% of samples. Table S3 summarizes the number of identified species in Chennai samples across these distinct groups based on their relative abundance.

**Figure 2:**
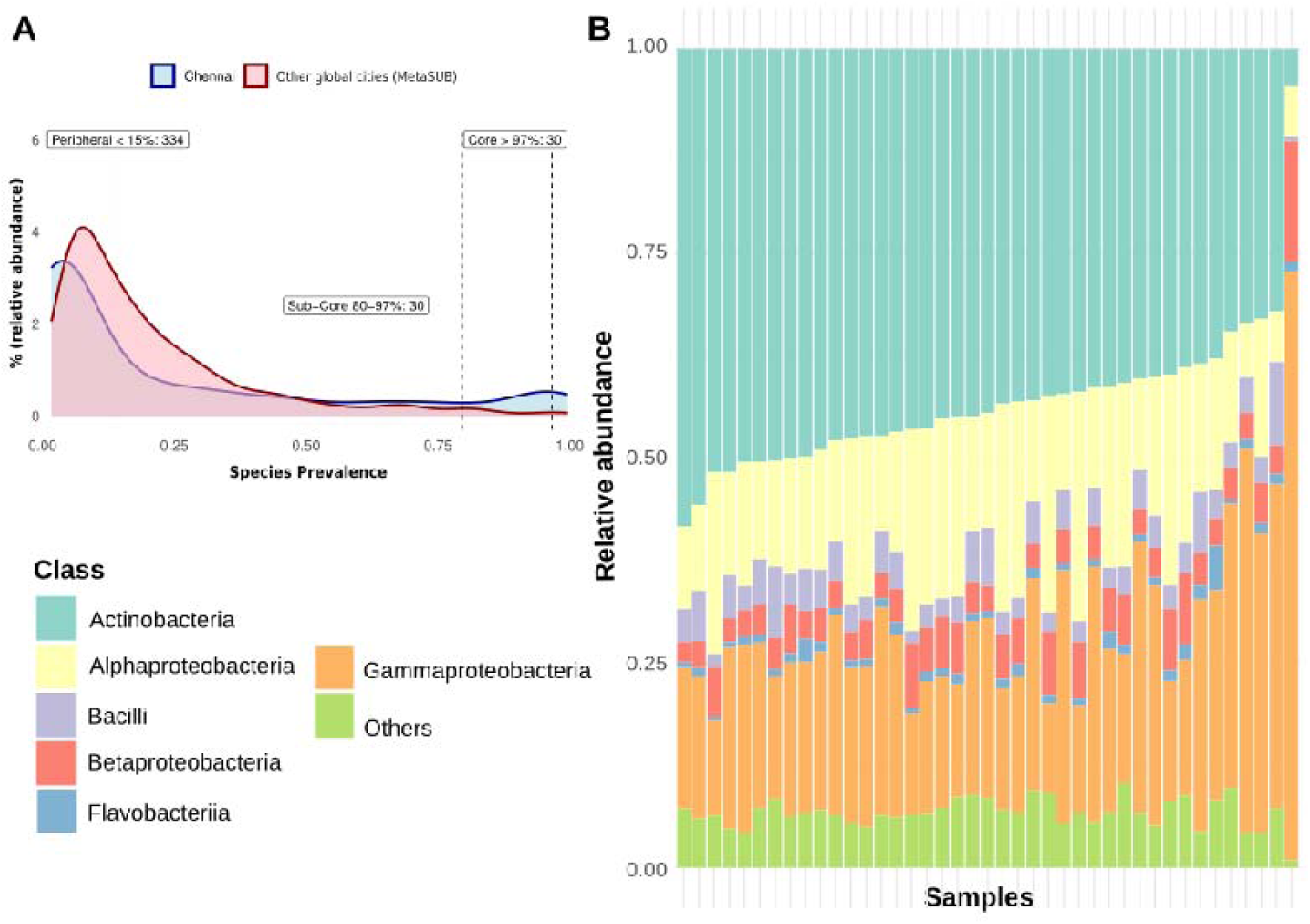
(A) Prevalence of species at relative abundance 0.001 and (B) Relative abundance of bacterial classes across 47 Chennai samples.

At a relative abundance of 0.001, which corresponded to more than 400 reads supporting a given species (based on the minimum library size shown in Table S2), we identified 33 core species, 30 sub-core species, and 334 peripheral species in the Chennai samples (Supplementary File 2, Sheet 1). Most samples were dominated by species from three classes – *Actinobacteria, Alphaproteobacteria,* and *Gammaproteobacteria,* although the relative abundance of these classes varied (Figure 2B).

Geographical regions and urban environments significantly influence the composition and diversity of microbial communities, shaped by factors such as climate, air quality, human activities, and the built environment [36]. To explore these effects, we compared samples from Chennai with those from 31 other global cities (Table S4), aiming to identify common and unique patterns in species distribution. We identified microbial signatures unique to Chennai. To emphasize these, we created a category called ‘unique-core,’ which included core species specific to Chennai that were not core or sub-core in any other city at the same relative abundance. We found nine such species that were part of the core in Chennai but not in other global cities (Table 1, Table S3).

**Table 1:** Eight species differentially prevalent in Chennai city and the habitats in which they are typically found. These species are selected based on a threshold of <80% prevalence at a relative abundance of >0.001 in other global cities, whereas they are >97% prevalent in Chennai at the same relative abundance.

| Species | Sources |
| --- | --- |
| *Brachybacterium sp. SGAir0954* | Isolated from tropical air collected in Singapore [48] |
| *Brachybacterium faecium* | Isolated from poultry deep litter [24] |
| *Brachybacterium saurashtrense* | Isolated from the roots of *Salicornia brachiata*, an extreme halophyte [23] |
| *Pseudomonas mendocina* | Found in water and soil samples [28] |
| *Pseudomonas alcaligenes* | Commonly found in soil and water [29] |
| *Staphylococcus arlettae* | Found in both environment and human sources [25–27] |
| *Sphingobium yanoikuyae* | Found in environments ranging from terrestrial to aqueous habitats, including seawater [30] |
| *Comamonas aquatica* | Found in water source [31] |
| *Brachybacterium sp. P6-10-X1* | Isolated from deep-sea sediments [49] |

Among these, *Brachybacterium saurashtrense*, which is adapted to high salinity, was first isolated from the roots of the halophytic plant *Salicornia brachiate* [23]. Its presence indicates that Chennai’s coastal climate and humidity support its growth. Similarly, *Brachybacterium sp. SGAir0954*, previously isolated from tropical air in Singapore [25], thrived in Chennai’s tropical and humid conditions. *Brachybacterium faecium* [24], typically associated with poultry environments, suggested agricultural activities near Chennai or the transportation of animal products. The presence of human-associated species, such as *Staphylococcus arlettae* [25–27], commonly found on the skin, highlighted the roles of metro passengers and urban density in microbial spread. Soil- and water-associated species, such as *Pseudomonas mendocina* [28], *Pseudomonas alcaligenes* [29], *Sphingobium yanoikuyae* [30], and *Comamonas aquatica* [31], likely reflected the local soil and water microbiome, demonstrating resilience to pollutants and potential for bioremediation in urban ecosystems. Overall, these findings illustrated how Chennai’s tropical climate, coastal characteristics, dense human activity, and proximity to agricultural areas shaped the unique microbiome of its metro stations.

### Microbial diversity and composition across various surface types

We evaluated the quantity and diversity of microbial communities across different surface types by analyzing alpha and beta diversity. We used the Shannon-Wiener species diversity index to assess alpha diversity, which accounts for both species richness (the number of species present) and species evenness (the distribution of individuals among species). The samples collected from the kiosk exhibited a significantly higher Shannon index (median = 6.73), indicating greater species diversity than on other surfaces. In contrast, the samples from the rods showed the lowest Shannon index (median = 5.51), suggesting lower diversity (Figure 3A). For beta diversity, we performed Principal Coordinate Analysis (PCoA) using the Bray–Curtis dissimilarity measure to assess differences in species composition among samples from various surfaces. This analysis considered both the presence or absence of species and their relative abundances. As expected, the kiosk samples, which showed the highest species diversity, clustered separately, indicating a distinct species composition compared to samples from other surfaces (Figure 3B).

**Figure 3:**
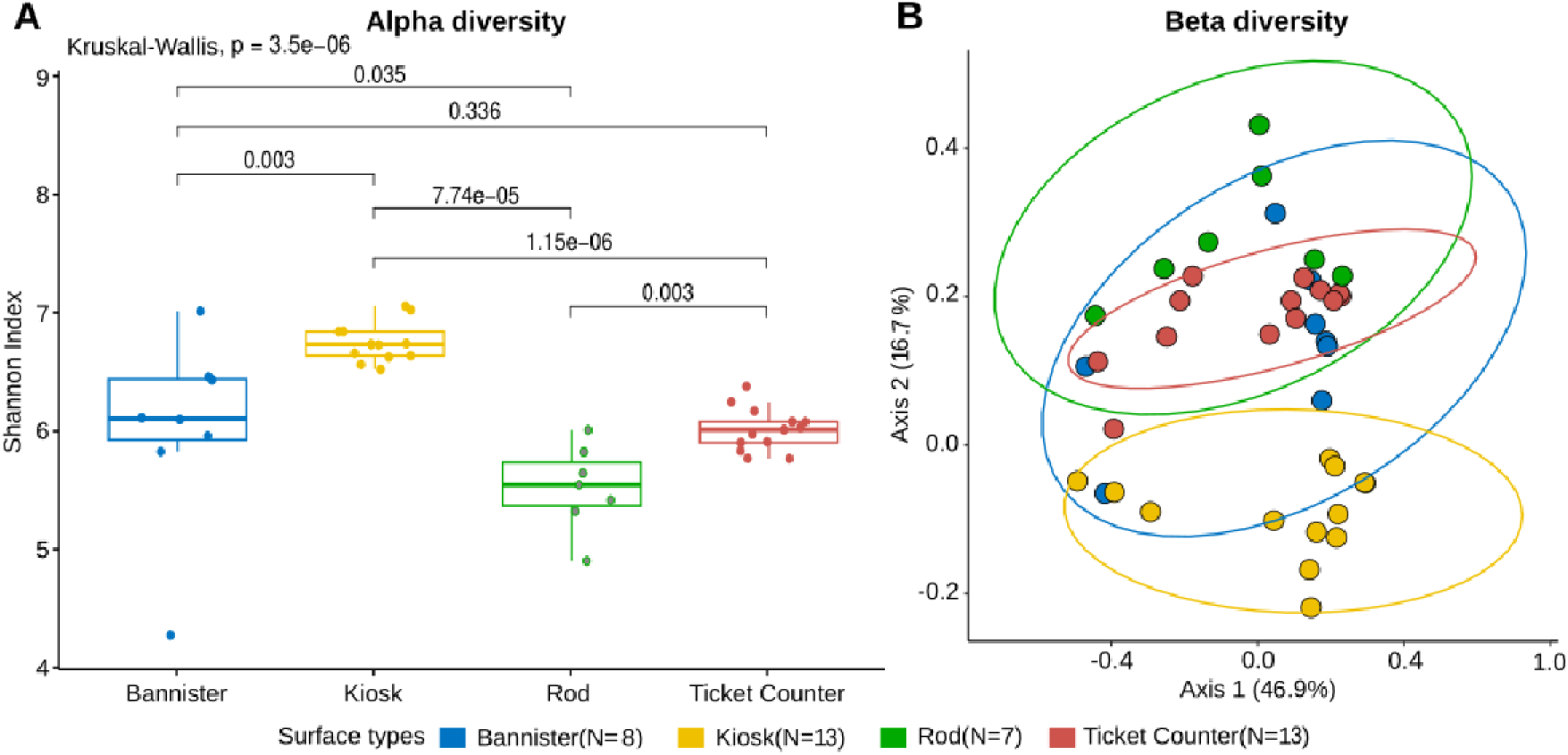
(A) Alpha diversity and (B) Beta diversity of Chennai samples collected from four different surface types. A “one-versus-rest” comparison was performed for each surface type in both analyses. For alpha diversity, Wilcoxon pairwise tests were used to compute p-values. For beta diversity, PERMANOVA was conducted to assess group-wise differences. No beta-diversity comparisons yielded p-values < 0.05.

We further examined the variations in microbial species across different surface types. We found that 19 species were significantly enriched on kiosk surfaces (|log_2_FC| > 1 and P_adj_ < 0.05) compared to all other surfaces, while five species were significantly enriched on the rod surfaces (Supplementary File 2, Sheet 5-6). This finding aligned with our diversity analysis, where kiosk surfaces, which exhibited greater species diversity, also had more species that were significantly overrepresented. In contrast, no species were significantly over- or underrepresented on the banister or ticket counter surfaces compared to the other surface types (Figure 4). The kiosk surfaces exhibited higher alpha diversity and a distinct microbial composition compared to other surfaces. Notably, several species, such as *Kocuria flava, Deinococcus sp. NW 56, Micropruina glycogenica, Rhodobacter sphaeroides*, and multiple

**Figure 4:**
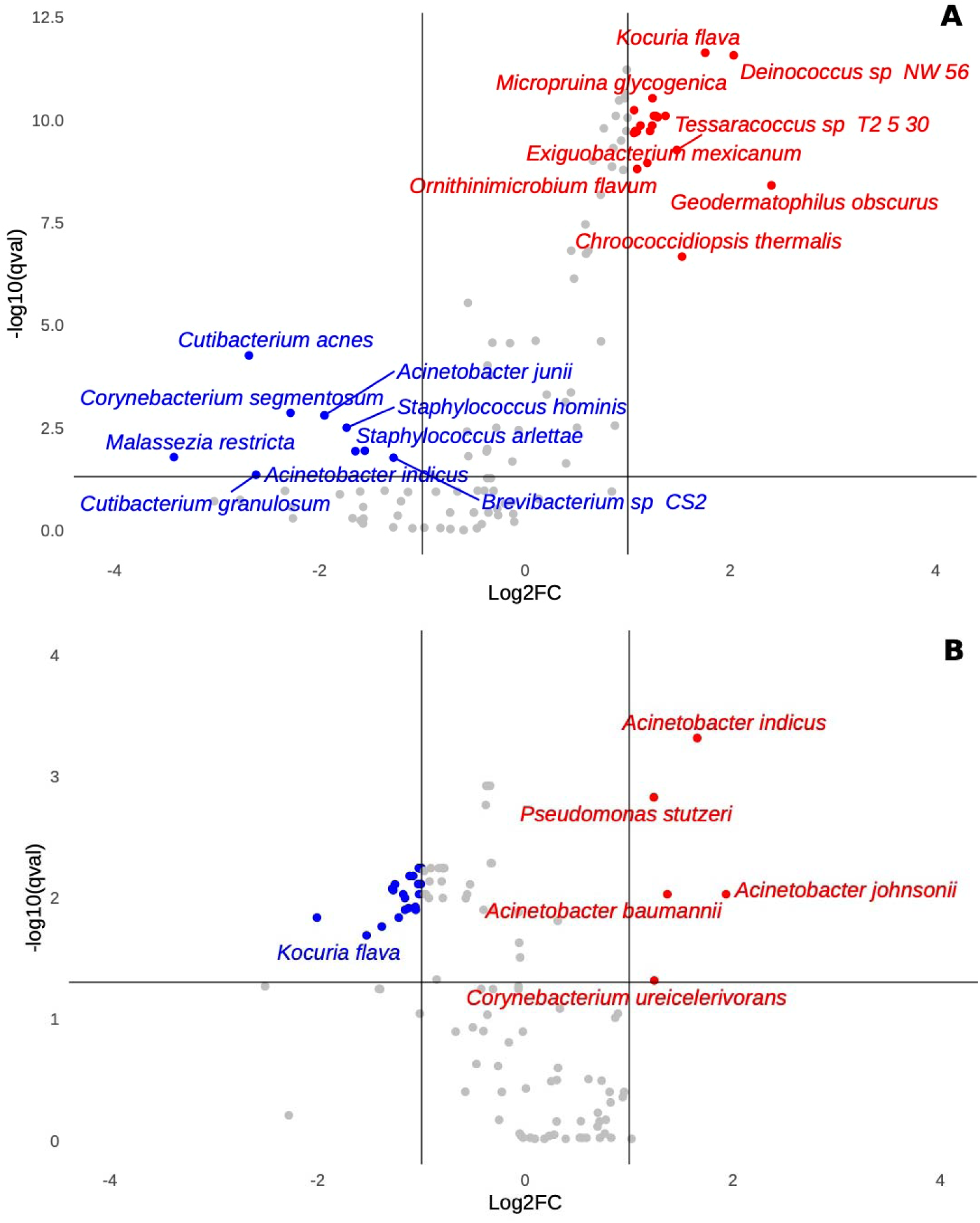
Differential representation of microbial species across surface types. (A) Comparison of species diversity on kiosk surfaces versus other surfaces. (B) Comparison of species diversity on rod surfaces versus other surfaces. Species with loglFC > 1 or loglFC < -1 and P_adj_ < 0.05 are shown in red and blue, respectively.

*Nocardioides spp*. were significantly overrepresented, indicating their predominance in this environment. In contrast, species like *Cutibacterium acnes*, *Corynebacterium segmentsum*, and *Staphylococcus hominis* were underrepresented, suggesting differential surface colonization patterns. The presence of opportunistic pathogens and resilient bacteria on high-contact kiosk surfaces underscores the importance of targeted cleaning protocols to mitigate potential health risks.

### Pangenome profiling of Metagenome-assembled Genomes (MAGs)

We investigated the genetic diversity and functional capabilities of microbial communities in the Chennai metro system through pangenome analysis of MAGs. We reconstructed 47 high-quality MAGs from the samples, each satisfying the standard threshold of >90% completeness and <5% contamination, making them suitable for downstream pangenome analysis. These MAGs were initially assigned at the genus level, resulting in the identification of 12 genera (Figure 5A). Of these, 44 MAGs spanning 10 genera could be further resolved to the species level (Supplementary File 2, Sheet 2). MAGs assigned to the genera *Brevilactibacter* and *Saccharomonospora* did not meet the criteria for confident species-level classification and were therefore reported as “Unknown species” in Figure 5A. We identified 11 species with average nucleotide identity (ANI) greater than 97% compared to reference genomes. The identified species included *Acinetobacter fasciculus, Acinetobacter junii, Brevibacterium paucivorans, Corynebacterium senegalense, Cutibacterium acnes, Epilithonimonas bovis, Kocuria marina, Micrococcus luteus, Moraxella osloensis, Stutzerimonas stutzeri,* and *Weissella confusa.* Among the identified species, *Cutibacterium acnes* was represented by 18 MAGs, *Stutzerimonas stutzeri* by 11 MAGs, and *Micrococcus luteus* by 7 MAGs, while the remaining species were represented by a single MAG each (Figure 5A; Supplementary File 2, Sheet 2).

**Figure 5:**
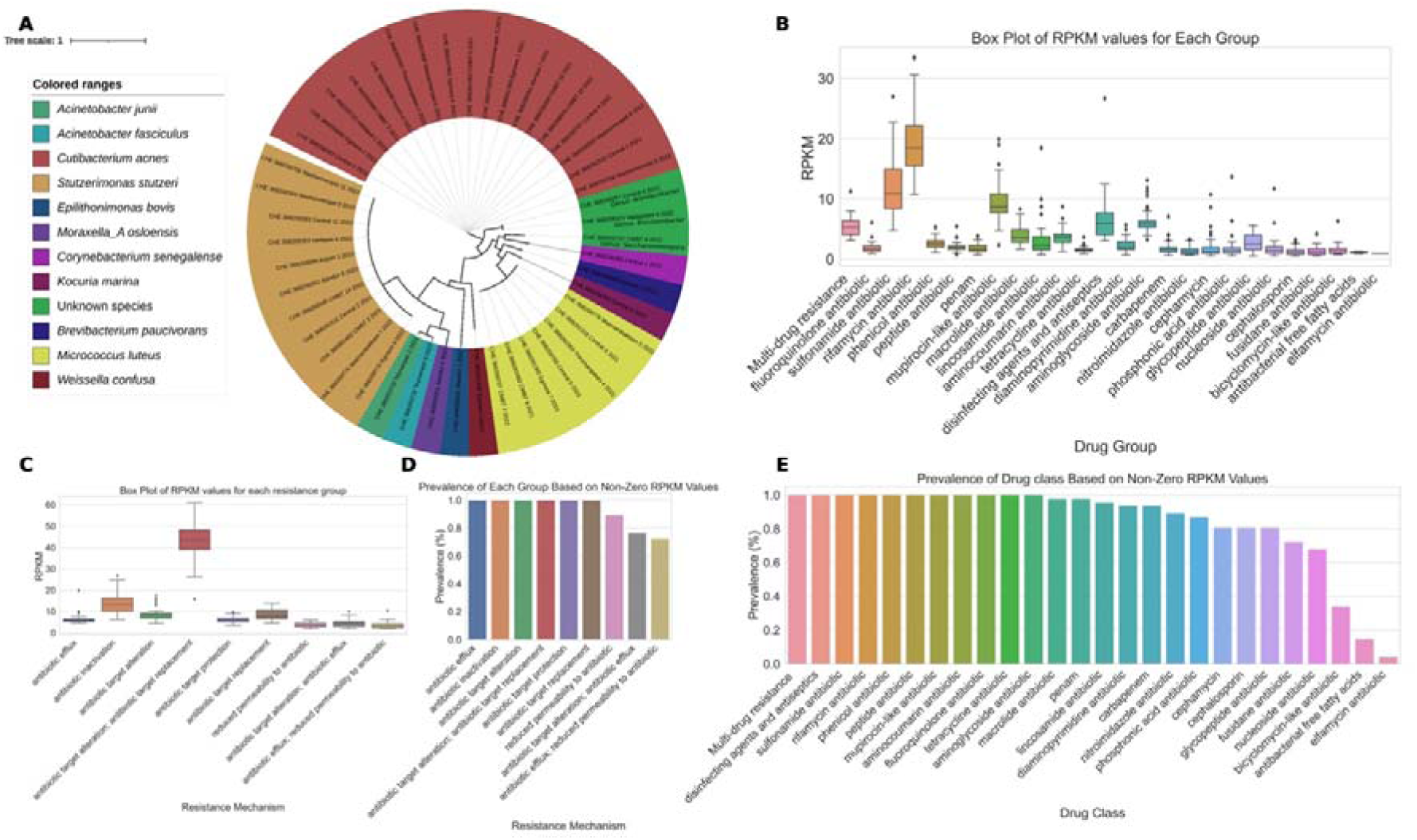
(A) Phylogenetic tree of the microbial species mapped to the MAGs. Gene abundance profiles and prevalence of AMR gene families across drug classes (B, D) and resistance mechanisms (C, E).

Our functional enrichment analysis using the COG2020 database identified a significant overrepresentation of certain functions in the MAGs from Chennai. Specifically, for *C. acnes*, we examined 18 MAGs from Chennai alongside 39 reference strains and found ten functions that were significantly overrepresented (Table 2). In the case of *S. stutzeri*, we analyzed 11 MAGs from Chennai and 20 reference strains, revealing four functionalities that were significantly overrepresented in the Chennai MAGs (Table 3). We did not identify any significant functional enrichment in *M. luteus* MAGs.

**Table 2:**
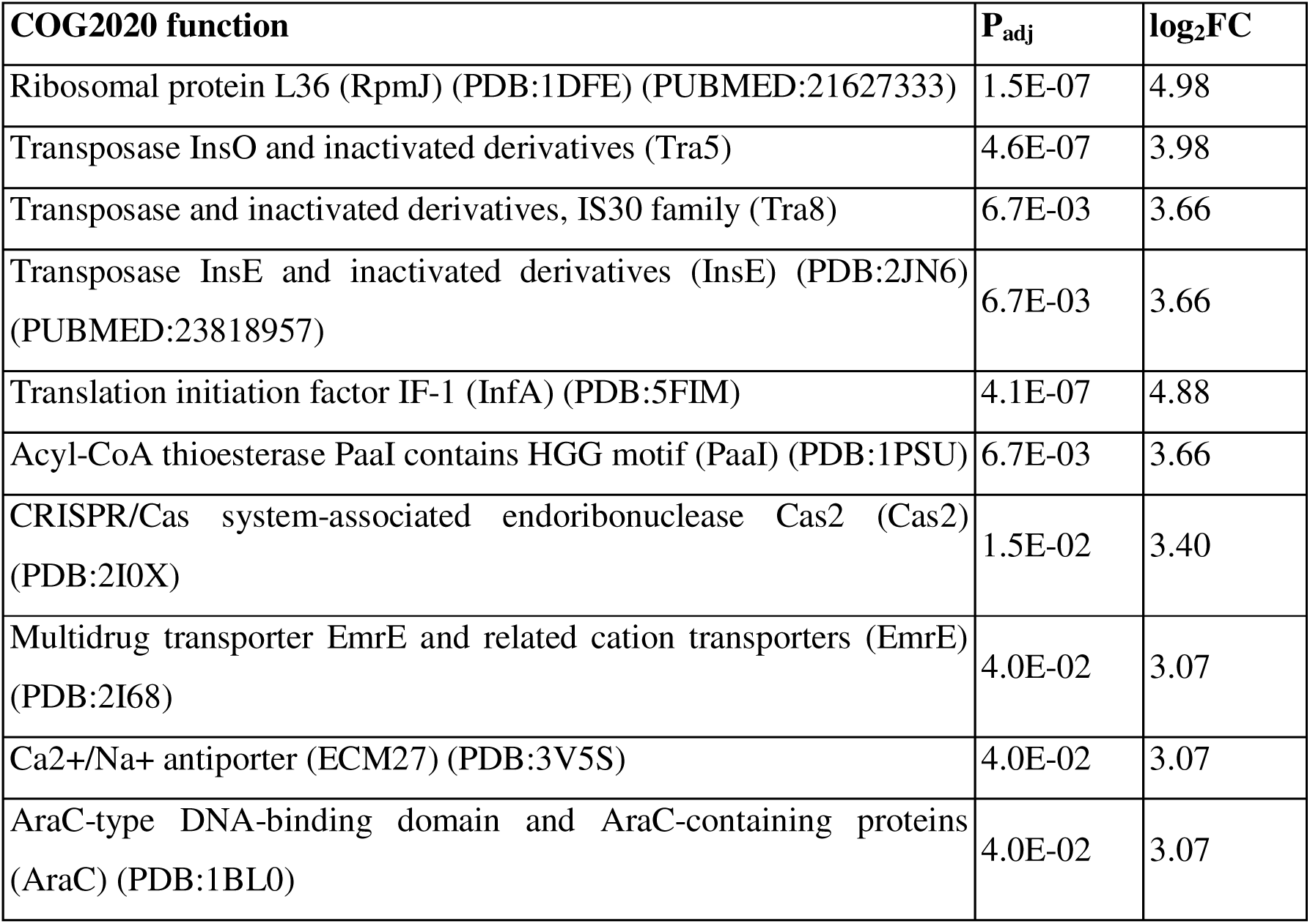
Significantly overrepresented functions in *Cutibacterium acnes* MAGs.

| COG2020 function | P<sub>adj</sub> | log<sub>2</sub>FC |
| --- | --- | --- |
| Ribosomal protein L36 (RpmJ) (PDB:1DFE) (PUBMED:21627333) | 1.5E-07 | 4.98 |
| Transposase InsO and inactivated derivatives (Tra5) | 4.6E-07 | 3.98 |
| Transposase and inactivated derivatives, IS30 family (Tra8) | 6.7E-03 | 3.66 |
| Transposase InsE and inactivated derivatives (InsE) (PDB:2JN6) (PUBMED:23818957) | 6.7E-03 | 3.66 |
| Translation initiation factor IF-1 (InfA) (PDB:5FIM) | 4.1E-07 | 4.88 |
| Acyl-CoA thioesterase PaaI contains HGG motif (PaaI) (PDB:1PSU) | 6.7E-03 | 3.66 |
| CRISPR/Cas system-associated endoribonuclease Cas2 (Cas2) (PDB:2I0X) | 1.5E-02 | 3.40 |
| Multidrug transporter EmrE and related cation transporters (EmrE) (PDB:2I68) | 4.0E-02 | 3.07 |
| Ca<sup>2+</sup>/Na<sup>+</sup> antiporter (ECM27) (PDB:3V5S) | 4.0E-02 | 3.07 |
| AraC-type DNA-binding domain and AraC-containing proteins (AraC) (PDB:1BL0) | 4.0E-02 | 3.07 |

**Table 3:**
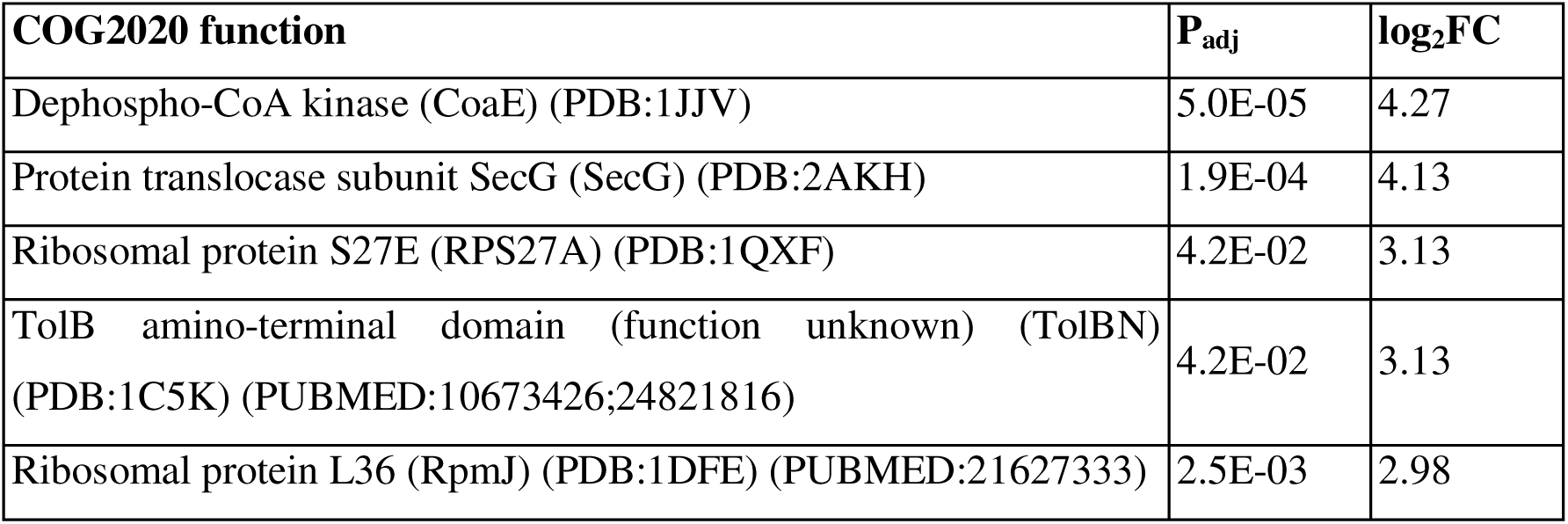
Significantly overrepresented functions in *Stutzerimonas stutzeri* MAGs.

The enrichment of multiple transposases among the *C. acnes* MAGs indicated that these MAGs possess mobile genetic elements that could facilitate genetic diversity and adaptability. These elements could help the organism acquire new traits or resistances, particularly in changing environments [32].

### Assessment of antibiotic resistance gene abundance

We annotated 206 AMR gene families in our sample set, associated with 25 drug classes and nine resistance mechanisms (Supplementary File 2; Sheet 3). We introduced a new class, “Multidrug resistance,” for genes that confer resistance to multiple antibiotic classes. Overall, the AMR gene families exhibited a low gene abundance pattern (Average RPKM – mean: 3.64, median: 1.13) across different drug classes and the resistance mechanisms (Supplementary File 2; Sheet 4). However, certain antibiotic classes, namely Sulfonamides and Rifamycins, had mean RPKM values of 12.29 and 19.25, respectively, indicating higher gene abundance than other drug classes (Figure 5B). According to the 2019 WHO AWaRe (Access, Watch, Reserve) classification framework [33], the Sulfonamide drug class is categorized as “Access,” while Rifamycin is classified as “Watch.” Interestingly, although the abundance of AMR classes was low, they were prevalent throughout our sample set, with only Bicyclomycin-like antibiotics, antimicrobial-free fatty acids, and Eifamycin antibiotics showing prevalence below 40% (Figure 5E).

Furthermore, the gene abundance pattern of the AMR gene family associated with the “antibiotic target alteration and antibiotic target replacement” resistance mechanism was notably higher than that of other mechanisms (Figure 5C). Overall, the prevalence of all AMR gene families corresponding to the annotated resistance mechanisms in our sample set exceeded 50% (Figure 5D).

## DISCUSSION

Urban public transit systems, such as metro networks, serve as critical hubs for microbial transmission, shaping the urban microbiome through constant human activity and high population density [34]. In densely populated regions like India, where millions rely on metro systems daily, these environments act as dynamic ecosystems where microbial communities interact, adapt, and spread. The Chennai Metro, operated by CMRL, has rapidly become a vital part of the city’s urban transit infrastructure. Since its launch on June 29, 2015, the metro has transported an impressive 355.3 million passengers, highlighting its critical role in urban mobility [35]. In 2024 alone, CMRL recorded 105.2 million passengers, a substantial increase from 91.1 million in 2023. As metro ridership continues to grow, so does the importance of understanding its impact on microbial transmission and urban health.

This study presents the first comprehensive metagenomic analysis of an Indian transit system, revealing a unique microbial signature with eight core species that were differentially abundant in Chennai. The presence of species like *Brachybacterium saurashtrense* and *Brachybacterium sp. SGAir0954* suggests a strong influence of Chennai’s coastal climate and humidity, potentially indicating marine spillover effects. Given Chennai’s location on the Coromandel Coast of the Bay of Bengal, this region has historically been affected by extreme weather events, such as the 2004 Indian Ocean tsunami [36]. Our environmental metagenomic findings also align with clinical evidence, highlighting region-specific challenges in antimicrobial resistance. For instance, a recent study in Vellore, Tamil Nadu, documented multidrug-resistant bacterial strains, including *Acinetobacter* species [37], in sewage receiving hospital wastewater, underscoring the selective enrichment driven by antibiotic residues and hospital effluents, which represent public health hazards through environmental dissemination. Moreover, locally prevalent resistant isolates displaying phenotypes relevant to rifamycin and sulfonamide resistance corroborate our AMR gene findings within the Chennai metro microbiome. These data reinforce the importance of including such environmental reservoirs in integrated AMR surveillance efforts.

Additionally, *Staphylococcus arlettae*, which we found abundantly represented in Chennai’s metro, is increasingly reported from clinical settings associated with medical device infections and bloodstream infections [25, 26], directly linking our environmental observations to the ecology of healthcare-associated pathogens. Opportunistic pathogens *Moraxella osloensis* and *Acinetobacter junii* are similarly known to persist in household and hospital environments within India [38], supporting their epidemiological relevance in urban microbial health risks. Therefore, monitoring these microbial communities in cities like Chennai is crucial.

The risk of increased flooding from sea-level rise [39] further underscores the need for continued surveillance. Flooding during monsoon seasons can disrupt urban microbial ecology by dispersing environmental microbes, chemical contaminants, and antibiotics into public spaces, such as metro stations. This process can lead to shifts in the resident microbial communities and enhance the spread of AMR genes, which may persist and proliferate in the post-flood environment. Such environmental perturbations pose a significant public health risk, particularly in densely populated areas with high human-microbe interactions. Therefore, regular microbial surveillance in flood-affected urban transit environments is crucial for detecting these dynamic changes early, enabling targeted interventions to prevent the dissemination of resistant strains and protect public health. Our dataset offers valuable insights into future shifts in urban microbiomes, underscoring the importance of continued research.

Surface type significantly influenced microbial diversity, with kiosks exhibiting the highest alpha diversity, consistent with previous studies [40], likely due to frequent human contact. The concentrated use of kiosks as focal points for user interaction by diverse individuals can create unique microbial communities, potentially resulting in higher alpha diversity despite their localized sampling areas, which are comparatively smaller surface contact points than handrails or banisters. In contrast, the larger surface areas of handrails may support a more homogeneous microbial community due to constant exposure to human contact, as explained by ecological principles. According to these principles, continuous and extensive surface exposure favors the dominance of well-adapted taxa, thereby decreasing community heterogeneity. Meanwhile, smaller surfaces, such as kiosks, although having less total area, receive diverse and frequent contacts that vary temporally and spatially, introducing a wider array of microbial inputs. The resulting heterogeneity of kiosk microhabitats supports higher alpha diversity and greater taxonomic variability. Understanding these dynamics is crucial for developing effective cleaning protocols and public health strategies in urban environments.

We investigated the genetic diversity and functional potential of microbial communities in the Chennai metro system by reconstructing 47 high-quality MAGs. Among these, *Cutibacterium acnes*, *Stutzerimonas stutzeri*, and *Micrococcus luteus* were the most frequently represented species, suggesting their ecological dominance or adaptation to the urban transit environment. In contrast, several species were represented by a single MAG, indicating their potential niche specialization or low abundance within the ecosystem. Notably, *Moraxella osloensis*, an opportunistic pathogen commonly found in household environments, has been previously reported as a dominant species on kitchen surfaces and sponges, with its prevalence influenced by cleaning methods [38]. Similarly, *Acinetobacter junii* is frequently identified in house microbiomes in India, highlighting its persistence in built environments [38]. Among *C. acnes* MAGs, we observed an enrichment of transposases, suggesting the presence of mobile genetic elements that could promote genetic diversity and adaptability [32]. These elements may facilitate the acquisition of new traits or resistance mechanisms, particularly under environmental stress. Additionally, the overrepresentation of the multidrug transporter *EmrE* in *C. acnes* implies a potential mechanism for exporting toxic compounds, including antibiotics, thereby enhancing its survival under antimicrobial pressures. In *S. stutzeri* MAGs, we observed an overrepresentation of Dephospho-CoA Kinase (*CoaE*), an enzyme critical for coenzyme A (CoA) biosynthesis [41]. The significant overrepresentation of *CoaE* underscores the bacterium’s metabolic adaptability and ecological versatility in urban transit environments. CoA biosynthesis is essential for fundamental cellular processes, including energy production, fatty acid metabolism, and responses to environmental stresses such as disinfectants and oxidative agents commonly encountered on public surfaces. *S. stutzeri* is a well-documented opportunistic pathogen with the ability to persist across diverse environments, including soil, water, and clinical settings, and it has been implicated in bloodstream and respiratory infections, especially in immunocompromised individuals [42]. The elevated *CoaE* activity may enhance *S. stutzeri’s* resilience and persistence on frequently touched surfaces, thereby increasing the potential for microbial transmission and colonization in crowded urban settings. Given that CoA biosynthesis enzymes also represent antimicrobial drug targets [43], this links the pathway to clinically relevant resistance mechanisms. This finding highlights a functional biomarker of microbial survival and public health risk, advocating for integrated microbial and metabolic surveillance in urban transit systems to inform cleaning and infection control strategies. Our AMR gene analysis revealed low overall abundance across most drug classes, yet their widespread presence indicates latent resistance potential. Genes associated with Sulfonamide and Rifamycin resistance exhibited relatively higher abundance, suggesting selective pressure. Sulfonamides act as broad-spectrum antibiotics by inhibiting dihydropteroate synthase [44], while Rifamycins target bacterial RNA polymerase [45]. Notably, the high prevalence of Rifamycin resistance aligns with reports of its presence in hospital isolates from Chennai, indicating possible overlap between clinical and urban microbial populations [46]. This selective pressure likely arises from environmental antibiotic exposure resulting from human healthcare, agriculture, and community use, compounded by the use of disinfectants that co-select resistance traits, enabling bacterial survival in urban transit settings. Sulfonamide resistance typically involves alternative dihydropteroate synthase enzymes, while rifamycin resistance arises primarily from mutations in the RNA polymerase β subunit, both of which reduce antibiotic efficacy. Notably, AMR genes were predominantly linked to “antibiotic target alteration” and “antibiotic target replacement” mechanisms, which modify or substitute molecular targets, such as penicillin-binding proteins (β-lactams) or ribosomal RNA (macrolides). This dominance suggests that target modification is a primary adaptive strategy in these metropolitan environments, shaped by selective pressure from common antibiotic classes.

Our findings underscore the importance of continuous microbial surveillance in urban public transportation systems, particularly in dense and rapidly growing cities like Chennai. As urbanization accelerates and public transport networks expand, the dynamics of microbial transmission in these densely populated environments will become increasingly relevant to public health. This aligns directly with the UN Sustainable Development Goals (SDGs), particularly SDG 3 (Good Health and Well-Being) and SDG 11 (Sustainable Cities and Communities), which emphasize the need for resilient urban infrastructure that promotes health and sustainability. Integrating microbial surveillance into public health strategies is essential for achieving these goals, as it enables real-time monitoring of potential risks, such as AMR and the spread of infectious diseases.

Considering the growing emphasis on building smart cities, where technology-driven solutions enhance urban living, we posit that real-time metagenomic analysis can be pivotal in making urban spaces healthier and more resilient. Smart cities are increasingly integrating IoT-based systems for air quality, traffic, and waste management, but microbial surveillance should also be a core component of these ecosystems. Future research should prioritize long-term monitoring of microbial communities in urban environments, enabling city planners to design more adaptive infrastructures that mitigate public health risks while promoting sustainable development.

The unique microbial signatures identified in Chennai’s Metro further emphasize the need for region-specific strategies to address microbial transmission in public spaces. Comparing urban microbiomes across global cities could help policymakers develop targeted cleaning protocols, optimize public health interventions, and manage AMR more effectively. Future research should investigate how human behaviour, environmental factors, and urban design impact microbial community composition, and should use this knowledge to develop more resilient, health-conscious urban infrastructure within the broader framework of sustainable and smart city development.

## METHODS

### Sampling microbial communities in the Chennai Metro stations

To explore microbial communities within the Chennai Metro system, we collected samples (n = 96) from 12 metro trains and stations along the Blue and Green Lines, operated by Chennai Metro Rail Limited (CMRL; Figure 1A). The Blue Line spans 32.65 km, connecting Wimco Nagar Depot to Chennai International Airport, while the Green Line covers 22 km from Chennai Central to St. Thomas. Sampling occurred in two batches: December 2021 and January 2022.

Using the MetaSUB protocol [12], samples were collected with individually packaged Isohelix Buccal Mini Swabs (MS Mini DNA/RNA Swab, Isohelix, Cat.: MS-02) and stored in barcoded 2D Matrix V-Bottom ScrewTop Tubes (Thermo Scientific, Cat.: 3741-WP1D-BR/1.0mL) pre-filled with 400 µL of Zymo Shield transport and storage solution (Zymo Research, Cat.: R1100) to preserve DNA and RNA. Immediately after collection, swabs were placed in the matrix tubes and stored at -80°C until DNA extraction.

The samples were taken from frequently touched surfaces, including bannisters (handrails inside metro stations), kiosks (ticket vending machines), rods (handrails inside trains), and ticket counters (Figure 1B). Four samples were collected per station, totaling 48 per batch. Additionally, one negative control (an air sample) was collected at the Chennai Airport metro station by holding a dampened Copan swab exposed to the air for approximately 3 minutes, following the MetaSUB consortium protocol [4]. This comprehensive sampling strategy targeted diverse interaction points in the transit environment, providing a robust representation of microbial diversity. Out of 96 collected samples, 47 passed quality control (QC) criteria and were subjected to shotgun metagenomic sequencing for downstream analysis (Figure 1C). Of 47 samples from Chennai, 6 had fewer than 1 million reads after preprocessing and were excluded from further analysis. The remaining samples had an average of 5.4 million reads. A detailed list of all collected samples is provided in Supplementary Table S1.

### DNA Extraction, Library Preparation, and Sequencing

DNA extraction was performed using the Microlab STAR Liquid Handling System (Hamilton, Cat.: Microlab STAR) and the ZymoBIOMICS 96 MagBead DNA Kit (Zymo Research, Cat.: D4308) to ensure high-quality recovery of both abundant and low-abundance microbial species. An initial QC was conducted using the Qubit dsDNA High-Sensitivity Assay. A minimum DNA concentration of 0.9 ng was required for samples to pass QC after extraction. Libraries were prepared using the KAPA HyperPrep Kit (Roche), and sequencing was performed on the Illumina NovaSeq 6000 platform, chosen for its ability to generate high-throughput, paired-end reads suitable for metagenomic analysis. These methods enabled us to maximize the coverage of the microbial diversity present in the samples. Sample processing, sample DNA extraction, and sequencing were performed by MedGenome Labs Ltd., Bangalore, India.

### Preprocessing of the raw reads and mapping

To characterize the bacterial communities present in Chennai’s metro stations, bacterial taxa were identified through DNA sequencing. The raw Illumina sequencing reads were first assessed for quality using FastQC v0.11.9, and MultiQC v1.13 was employed to aggregate and visualize the quality reports. Preprocessing and assembly of the reads were conducted using modules from the MetaSUB Core Modular Analysis Pipeline (CAMP; https://github.com/MetaSUB-CAMP) [13]. The preprocessing steps included filtering out low-quality bases, low-complexity regions, and short reads using fastp v0.22.0. Adapter sequences were removed from the filtered reads with AdapterRemoval v2.3.3. Error correction was performed on the filtered reads using the Tadpole tool from the BBTools v39.01 aligner package. Tadpole corrected sequencing errors by comparing overlapping reads, identifying mismatches and gaps, and generating a consensus sequence based on the majority of bases. This process reduced noise and improved accuracy by correcting base-calling errors. After preprocessing, the quality of the corrected reads was reassessed using FastQC and MultiQC to ensure the integrity of the dataset. Finally, the processed reads were mapped to host reference genomes, including the human genome assembly (GRCh38) and the mouse genome assembly (MM39), using Bowtie2 v2.4.5 with the ‘very-sensitive’ flag (Table S2). Reads mapped to the host genomes were removed to focus on microbial sequences. After preprocessing, we selected 41 samples for analysis that had >1 million reads.

### Taxonomic classification and identification of the core and sub-core species

For taxonomic classification and determination of species relative abundance across all metagenomic samples, we used Kraken2 v2.1.3 and Bracken v2.7 from the CAMP pipeline. Kraken2 matched individual k-mers within query sequences to the lowest common ancestor (LCA) among all genomes containing the respective k-mer in the kraken2 database v0.1.1 made available by CAMP. To estimate species-level relative abundance, we used Bracken v2.7, which probabilistically redistributed reads from higher taxonomic levels to refine abundance estimates. We defined the core and sub-core species based on the distribution of taxa prevalence across our dataset. Here, prevalence refers to the fraction of samples in which a particular taxon was found at any abundance. This combined Kraken2-Bracken approach was chosen for its ability to balance computational efficiency with taxonomic depth, a critical feature for analyzing large, diverse datasets like those generated in this study. By leveraging these tools, we ensured both speed and accuracy in taxonomic classification, which was essential for subsequent comparative analyses with global urban microbiome datasets.

### Comparative analysis against the global urban microbiome composition

To compare the microbiome profile of Chennai with those of other major global cities, we utilized data from the global metagenomic map of urban microbiomes provided by the MetaSUB consortium [4]. We performed a batch download of clean read FASTQ files from the MetaSUB central data repository using Geoseeq v0.2.8 [14]. These clean reads were pre-processed through the CAP2 pipeline, which included filtering out low-quality bases and removing host reads. In total, 2,515 samples from 38 cities were initially obtained based on data availability at the time of download. Using the associated metadata for each sample, we excluded those originating from air or biological sources, focusing only on surface samples. This filtering step yielded a final dataset comprising 2,461 samples from 31 cities (Supplementary File 2, Sheet 7). The city with the fewest samples contributed 12, and the total number of samples used per city in our analysis is listed in Table S4. We conducted taxonomic classification analysis on the selected samples using Kraken2 and Bracken as implemented in the CAMP pipeline, consistent with our approach for the Chennai city samples. This allowed us to classify detected species in the Chennai dataset into core, sub-core, peripheral, and unique-core categories across different relative abundance thresholds (Table S3). Subsequently, we compared these species across all cities, including Chennai, to detect any unique signatures specific to Chennai and to identify the sample origin locations.

### Diversity analysis

To assess species richness (the variety of species) and evenness (their relative abundance), providing insights into microbial ecosystem differences across environments, we conducted diversity analyses using the phyloseq v1.44.0 [15] package in R. The phyloseq package integrates with the vegan package for calculating alpha diversity metrics, while ggpubr v0.6.0 [16] was used for visualization. Alpha Diversity (within-sample diversity) was quantified using the Shannon Index, which measures species distribution entropy. Higher values indicate greater diversity by accounting for both richness and evenness. We used the Wilcoxon test for pairwise comparisons and the Kruskal-Wallis test for group comparisons. The p-values from the pairwise comparisons were adjusted (P_adj_) using the Benjamini-Hochberg (BH) method in R. Beta Diversity (between-sample diversity) was analyzed using the “ordinate” function in phyloseq, applying Bray–Curtis dissimilarity and Principal Coordinate Analysis (PCoA). Bray–Curtis dissimilarity accounts for taxonomic abundance, providing a sensitive measure for detecting differences between samples. “One-versus-rest” comparison was performed for each of the four sample collection surfaces using the PERMANOVA test.

### Differential taxonomic profiles across sample collection objects

To examine variations in microbial community composition across sample collection objects, we performed a differential abundance analysis using MaAsLin2 v1.15.1 [17]. We employed a “one-versus-rest” approach, comparing each surface type (kiosks, rods, bannisters, ticket counters) against all other surfaces combined to identify taxa significantly enriched on each surface. Specifically, for kiosks, we conducted separate comparisons against rods, bannisters, and ticket counters. MaAsLin2 applied a generalized linear model to relative abundance data, with surface type as a fixed effect. Taxa with an absolute log-fold change |LFC| > 1 and a P_adj_ < 0.05 were considered significantly differentially abundant. This method enabled us to characterize and quantify microbial differences associated with surface types, highlighting key taxa that are influenced by the collection environment.

### Metagenomic assembly and genome reconstruction

To reconstruct microbial genomes from the metagenomic data, we first assembled the processed sequencing reads using MetaSPAdes v3.15.5 [17]. This produced a collection of contigs and scaffolds representing genomic fragments recovered from the samples. Following the assembly, MetaBAT2 v2.2.7 [18] was employed to bin metagenome-assembled genomes (MAGs). MetaBAT2 used two primary metrics to accurately cluster contigs—coverage depth per contig and tetranucleotide frequency (TNF) patterns. Coverage depth reflected the frequency of nucleotide sequencing, which was determined by aligning raw reads to the assembled contigs. TNF patterns, which represent the frequency of all possible four-nucleotide sequences within a contig, helped distinguish contigs from different microbial genomes based on their genomic composition. To evaluate the quality of the reconstructed MAGs, we used CheckM v1.0.1 [19]. This tool assesses genome completeness and contamination by analyzing lineage-specific marker genes, providing crucial insights into the reliability of the MAGs for downstream biological inferences.

### Pangenome analysis

To explore genetic diversity within the microbial population, we conducted a pangenome analysis that compared multiple genomes to identify core genes shared across genomes and variable genes unique to individual genomes or subsets of genomes. We selected MAGs with completeness greater than 90% and contamination levels below 5%. Taxonomic annotations of the selected MAGs were performed using GTDB-Tk v2.1.1 [20], and reference genomes for identified organisms were retrieved from NCBI. We used Anvi’o v8 [17], a tool that facilitates visualization, gene and functional annotation, and enrichment analysis across genomes using the Clusters of Orthologous Groups (COG) database (COG2020 [21]). We additionally constructed a phylogenomic tree using GToTree v1.8.4 [17]. This tree was based on 15 single-copy gene Hidden Markov Models (HMMs) derived from the Pfam database and NCBI, providing insights into the evolutionary relationships among the genomes under study.

### Assessing the gene abundance profile of the AMR gene family

To assess the antibiotic resistance potential of the microbial communities present in the metro stations, we generated the antibiotic resistance profile of the samples using the Resistome Gene Identifier (RGI) tool v6.0.2 from the Comprehensive Antibiotic Resistance Database (CARD) [22] database v3.2.6. RGI-bwt function from the RGI tool aligned the short read to CARD’s protein homolog models using the k-mer alignment (KMA) read aligner and provided a table of identified drug classes, the annotation for the drug resistance mechanisms, AMR gene family, the number of reads mapped, and the coverage. The read counts from RGI-bwt were normalized by total reads per sample and gene length to calculate RPKM values, and the abundance levels of AMR genes across different antibiotic classes were estimated using CARD. We calculated the prevalence of AMR gene families in our sample set by identifying those present across various drug classes and resistance mechanisms. To reduce potential noise and focus on gene abundance, we put a threshold at the 25th percentile (Q1) of RPKM. For a given drug class or drug resistance mechanism, the prevalence was 100% if at least one AMR gene family was present in all samples. Further, we also used the 2019 WHO AWaRe (Access, Watch, Reserve) classification framework to categorize antibiotic classes associated with the identified AMR genes.

## LIST OF ABBREVIATIONS

AMR: Antimicrobial Resistance
ANI: Average Nucleotide Identity
AWaRe: Access, Watch, Reserve (WHO classification framework for antibiotics)
BBTools: Bioinformatics tools for sequence analysis
CARD: Comprehensive Antibiotic Resistance Database
CAP2: Core Analysis Pipeline 2
CAMP: Core Modular Analysis Pipeline
CMRL: Chennai Metro Rail Limited
COG2020: Clusters of Orthologous Genes 2020 database
HMMs: Hidden Markov Models
KMA: K-mer Alignment
MAGs: Metagenome-Assembled Genomes
MetaSUB: Metagenomics and Metadesign of Subways and Urban Biomes Consortium
NGS: Next-Generation Sequencing
PCoA: Principal Coordinate Analysis
QC: Quality Control
RPKM: Reads Per Kilobase of transcript per Million mapped reads
RGI: Resistome Gene Identifier
SRA: Sequence Read Archive (data repository)

## DECLARATIONS

### Ethics approval and consent to participate

Not applicable

### Consent for publication

Not applicable

### Availability of data and materials

The raw reads are available in the SRA under accession PRJNA1201944 at https://dataview.ncbi.nlm.nih.gov/object/PRJNA1201944?reviewer=mg6622losta4rhp87h4rtl40hq. The codes used in this paper are available in the GitHub repository at https://github.com/IBSE-IITM/MetaSUB-CMRL. The rest of the data supporting this article is provided within the article and its online supplementary materials.

## Funding

The authors acknowledge funding from the Robert Bosch Centre for Data Science and AI (SB/20-21/0602/BTRBCX/008481) to K.R. and H.S., and from the Wadhwani School of Data Science and AI, IIT Madras.

### Authors’ contributions

K.R., H.S., and B.P. conceived the study; K.R. and H.S. acquired funding and supervised the study; V.Y.S. and V.P.G. pre-processed the data and established the analysis pipeline; V.Y.S., V.P.G., S.S., R.L., and S.V. collected samples and performed the analysis; V.P.G., V.Y.S., and S.S. wrote the first draft of the manuscript; V.P.G., V.Y.S., S.S., K.R., and H.S. edited the manuscript; all authors reviewed and commented on the manuscript.

## Acknowledgments

We thank Chennai Metro Rail Limited (CMRL) and its employees at the respective stations for allowing us to collect samples from Chennai Metro stations and trains for our research on the city’s microbial landscape. We also sincerely thank Mr. Santhanam Moorthyraj from the Dhan Foundation for facilitating our connection with CMRL. We thank Christopher Mason, David Danko, Braden Tierney, and Krista Ryon, Weill Medical College of Cornell University, New York, for their advice, support in planning, and for providing kits for sample collection.

