## Supplementary File 1 for "Microbial communities on station and train surfaces in Chennai Metro: Insights into urban transit microbiome"

Table S1: Number of samples sequenced per metro station

| **S.No** | **Metro station** | **No of samples**  **Dec-2021** | **No of samples**  **Jan-2022** |
| --- | --- | --- | --- |
| 1 | Airport | 0 | 2 |
| 2 | Meenambakkam | 0 | 1 |
| 3 | Alandur | 1 | 2 |
| 4 | Guindy | 0 | 1 |
| 5 | Saidapet | 0 | 2 |
| 6 | Teynampet | 0 | 3 |
| 7 | Central Station | 3 | 4 |
| 8 | Washermanpet | 2 | 4 |
| 9 | Egmore | 2 | 2 |
| 10 | Thirumangalam | 2 | 2 |
| 11 | CMBT | 3 | 3 |
| 12 | Vadapalani | 1 | 1 |
| 13 | Negative control | 0 | 0 |
| 14 | **Total** | **14** | **27** |

Table S2: Total reads per sample after removing human and mouse reads

| **S.No** | **Sample ID** | **Number of reads** |
| --- | --- | --- |
| 1 | 368257968 | 61,37,371 |
| 2 | 368258054 | 17,94,329 |
| 3 | 368259600 | 23,97,440 |
| 4 | 368259989 | 64,86,810 |
| 5 | 368273379 | 37,30,160 |
| 6 | 368281318 | 28,51,412 |
| 7 | 368281322 | 14,96,154 |
| 8 | 368281337 | 12,25,754 |
| 9 | 368281369 | 54,57,282 |
| 10 | 368281482 | 51,12,232 |
| 11 | 368281490 | 73,14,928 |
| 12 | 368281495 | 63,07,599 |
| 13 | 368281511 | 1,01,32,285 |
| 14 | 368281532 | 25,55,890 |
| 15 | 368246110 | 89,43,782 |
| 16 | 368253732 | 1,15,30,804 |
| 17 | 368258345 | 74,38,425 |
| 18 | 368258351 | 57,23,807 |
| 19 | 368258353 | 67,44,363 |
| 20 | 368258369 | 68,78,396 |
| 21 | 368258383 | 72,75,716 |
| 22 | 368258393 | 66,10,649 |
| 23 | 368258417 | 44,16,381 |
| 24 | 368258423 | 71,63,138 |
| 25 | 368258431 | 77,06,800 |
| 26 | 368259698 | 33,34,353 |
| 27 | 368259701 | 21,56,282 |
| 28 | 368259706 | 75,59,655 |
| 29 | 368259707 | 61,14,749 |
| 30 | 368259710 | 11,06,067 |
| 31 | 368259718 | 16,56,915 |
| 32 | 368259737 | 44,07,434 |
| 33 | 368259760 | 66,75,378 |
| 34 | 368259764 | 13,90,679 |
| 35 | 368259770 | 72,56,608 |
| 36 | 368259776 | 75,24,716 |
| 37 | 368260393 | 63,85,754 |
| 38 | 368297838 | 12,32,670 |
| 39 | 368297861 | 67,20,967 |
| 40 | 368299257 | 99,11,130 |
| 41 | 368299282 | 62,76,148 |

Table S3: Number of detected species across different relative abundance thresholds (scale 0–1) within each species category (core, sub-core, peripheral, and unique-core) in the Chennai dataset.

| **Species Category** | **Relative abundance** | | | |
| --- | --- | --- | --- | --- |
|  | **> 0.1** | **> 0.01** | **> 0.001** | **> 0.0001** |
| Core | 0 | 2 | 30 | 495 |
| Sub-core | 0 | 1 | 30 | 481 |
| Peripheral | 5 | 47 | 334 | 823 |
| Unique-core | 0 | 0 | 9 | 16 |

Table S4: Number of samples used per Cities from MetaSUB

| **S.No.** | **City** | **Number of samples** |
| --- | --- | --- |
| 1 | Hong Kong | 652 |
| 2 | London | 421 |
| 3 | Ilorin | 195 |
| 4 | Singapore | 180 |
| 5 | New York City | 148 |
| 6 | Porto | 111 |
| 7 | Barcelona | 96 |
| 8 | Kyiv | 74 |
| 9 | Doha | 66 |
| 10 | Seoul | 60 |
| 11 | Zurich | 52 |
| 12 | Lisbon | 51 |
| 13 | Stockholm | 43 |
| 14 | Denver | 37 |
| 15 | Berlin | 36 |
| 16 | Offa | 17 |
| 17 | Santiago | 17 |
| 18 | Naples | 16 |
| 19 | Paris | 16 |
| 20 | Sacramento | 16 |
| 21 | Hamilton | 16 |
| 22 | Marseille | 16 |
| 23 | Sofia | 16 |
| 24 | Vienna | 15 |
| 25 | Sendai | 14 |
| 26 | Brisbane | 14 |
| 27 | Kuala Lumpur | 14 |
| 28 | Hanoi | 14 |
| 29 | Auckland | 13 |
| 30 | Minneapolis | 13 |
| 31 | Baltimore | 12 |
